# Three-dimensional organization of transzonal projections and other cytoplasmic extensions in mouse ovarian follicles

**DOI:** 10.1101/351262

**Authors:** Valentina Baena, Mark Terasaki

**Author notes:** Corresponding author: Mark Terasaki.

## Abstract

Each mammalian oocyte is nurtured by its own multi-cellular structure, the ovarian follicle. We used new methods for serial section electron microscopy to examine entire cells and their projections in mouse antral ovarian follicles. It is already known that cumulus cells send towards the oocyte thin cytoplasmic projections called transzonal projections (TZPs), which are crucial for normal oocyte development. We found that most TZPs do not reach the oocyte, and that they often branch and make gap junctions with each other. Furthermore, the connected TZPs are usually contacted on their shaft by oocyte microvilli. Mural granulosa cells were found to possess randomly oriented cytoplasmic projections that are strikingly similar to free-ended TZPs. We propose that granulosa cells use cytoplasmic projections to search for the oocyte, and cumulus cell differentiation results from a contact-mediated paracrine interaction with the oocyte.

## Introduction

The mammalian ovarian follicle is a complex tissue structure that nurtures the growth of the oocyte and also serves as the endocrine organ which supplies the female hormones estrogen and progesterone (Hawkins and Matzuk, 2008). In the large antral stage, a basal lamina encloses about 1000 granulosa cells, which form multiple layers around the oocyte. The 2-3 layers of cells adjacent to the oocyte are known as cumulus cells (or cumulus granulosa cells), while the cells in the outer layers of the follicle are known as mural granulosa cells.

The follicle begins development as a small oocyte surrounded by a single layer of thin somatic cells (“primordial follicle”) and grows to full size over the course of 3-4 estrus cycles (each cycle is ~4 days) (Hirshfield, 1991). Follicle development involves multiple paracrine interactions (Edson et al., 2009; Richards and Pangas, 2010). For instance, growth-differentiation factor-9 (GDF9) is synthesized by the oocyte and is required for the follicle to develop past the single layer stage (Dong et al., 1996).

Early follicle growth is autonomous but later, the follicle becomes responsive to follicle stimulating hormone (FSH) from the pituitary. This hormone stimulates the differentiation of cumulus cells and outer mural granulosa cells, as well as the final stages of growth. The mural granulosa cells synthesize estrogen, and the hypothalamus monitors the number of mural granulosa cells by sensing the estrogen present in the blood. When this reaches a threshold level, the hypothalamus signals to the pituitary to release a pulse of luteinizing hormone (LH) (Knobil, 1974). LH acts on the follicle to start the ovulation process: the mural granulosa cells are reprogrammed to synthesize progesterone, the oocyte resumes meiosis, and the cumulus cells reorganize (cumulus expansion) to be expelled from the follicle along with the oocyte.

Gap junctions connect all cells in the follicle and have a critical role in follicle development and function (Simon et al., 1997). Gap junctions transmit nutrients taken up by the granulosa cells to the oocyte (Sugiura et al. 2005). Furthermore, they transmit the LH signal throughout the follicle. The LH receptors are present only on the outer mural granulosa cells (Bortolussi et al., 1979). LH binding causes a reduction of cGMP in these cells, which in turn lowers the cGMP levels in other granulosa cells and in the oocyte by diffusing through the gap junctions (Shuhaibar et al., 2015). Elevated cGMP levels in the oocyte maintain it arrested in meiotic prophase, and the reduction of cGMP caused by LH reinitiates meiosis in preparation for fertilization (Norris et al., 2009). A parallel pathway involving EGF also lowers cGMP (Liu et al., 2014).

The gap junctions between cumulus cells and the oocyte are present on remarkable structures called transzonal projections (TZPs) (Li and Albertini, 2013; Clarke, 2018). These are thin cytoplasmic projections that originate from the cumulus cells and traverse the 3-5 micron-thick extracellular matrix of the oocyte (zona pellucida). They contact the oocyte surface at adherens junctions (Mora et al., 2012) and gap junctions (Gilula et al., 1978). Because TZPs provide the site of gap junctional communication and possibly of other interactions between the oocyte and cumulus cells, they are crucial structural elements of the follicle. However, due to their density and complexity, the three-dimensional organization of oocyte components, TZPs, and their junctions is poorly understood.

Automation and new computer capabilities have improved serial section electron microscopy so that it is much more feasible to produce large, three-dimensional fields of view at high resolution (Denk and Horstmann, 2004). We used the ATUM (automated tape-collecting ultramicrotome) method with scanning electron microscopy (SEM) (Kasthuri et al., 2015) to examine entire cells and their relationships to neighboring cells in mouse antral ovarian follicles. The images provide new information on follicle structure, particularly the inter-relationships of the TZPs and the oocyte cell surface. They also reveal the presence of cytoplasmic projections among granulosa cells within the follicle. As in other tissues and systems, these projections may be essential for cell communication during normal development and function (Ramirez-Weber and Kornberg, 1999; Inaba et al., 2015; Heimsath et al., 2017).

## Results

### Cumulus cells extend connected and free-ended transzonal projections (TZPs)

The observations to be described below were obtained from antral follicles from prepubertal mice without exogenous hormonal stimulation. At this stage, the oocyte is surrounded by a several micron-thick extracellular matrix, the zona pellucida. Cumulus cells on the other side of the zona pellucida contact the oocyte by sending cytoplasmic projections through the zona pellucida (TZPs) (Figure 1A).

**Figure 1.**
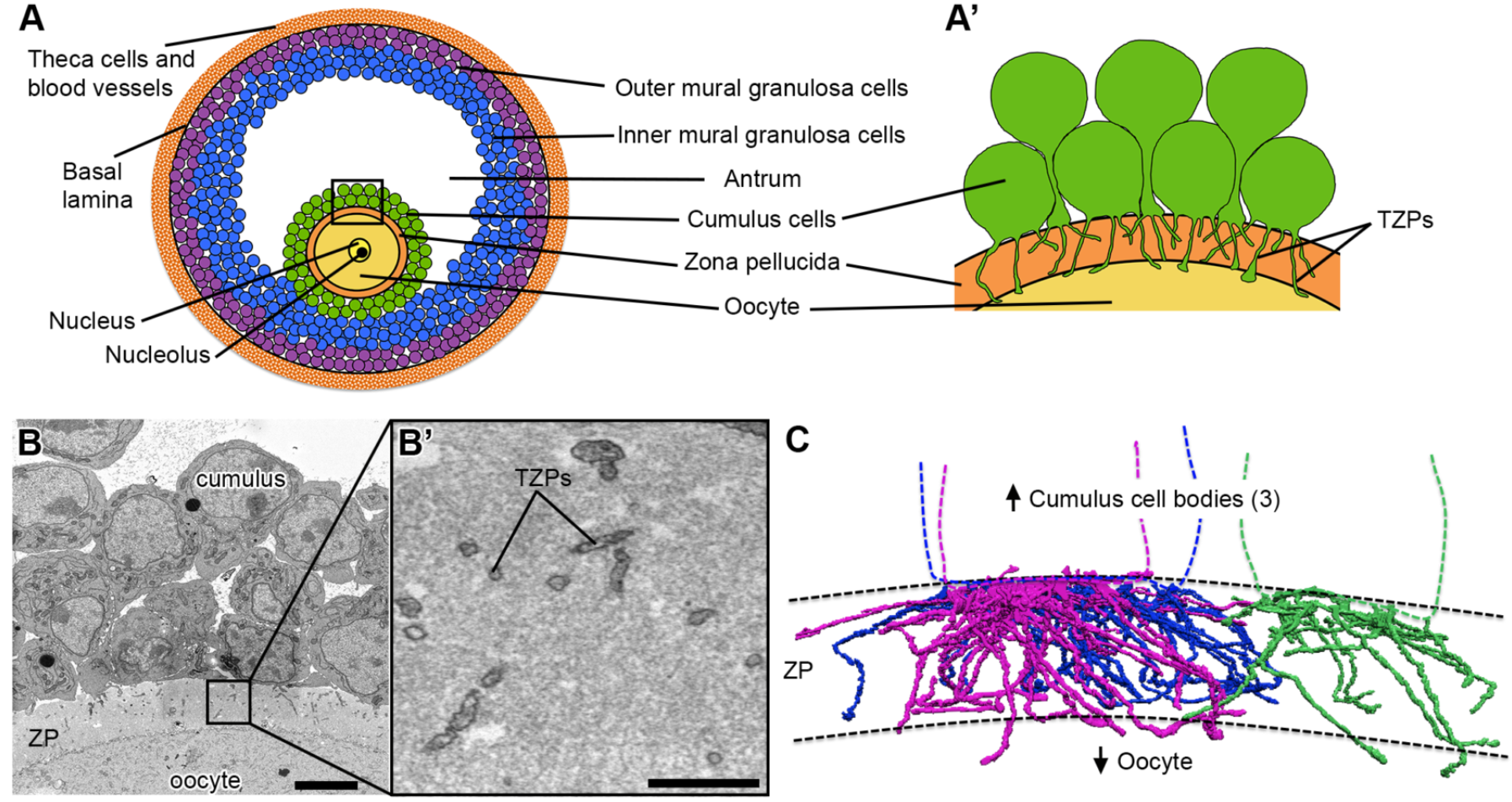
Cumulus cells send numerous projections through the zona pellucida. **(A)** Schematic of a mouse antral follicle. Black square represents area magnified in (A’). **(A’)** Schematic of several cumulus cells that send transzonal projections (TZPs) through the zona pellucida to connect to the oocyte. Cumulus cell bodies could be adjacent to or displaced from the zona pellucida. **(B)** SEM image of a cross-section through cumulus cells, zona pellucida (ZP), and oocyte of an antral follicle. Video 1 shows 202 serial sections of this area and highlights a cumulus cell with all of its projections. Scale bar, 5 μm. **(B’)** High-magnification of a region in the zona pellucida showing several TZPs in cross-section imaged by SEM. Scale bar, 1 μm. **(C)** Reconstruction of every TZP sent by three neighboring cumulus cells, each shown in a different color. TZPs were segmented from 405 serial electron micrographs, encompassing a volume of 22.5 × 6.7 × 18.2 μm (x, y, z). ZP, zona pellucida.

To visualize TZPs in three dimensions, we collected ~600 serial sections of 40-45 nm thickness (total depth 27 μm) and imaged volumes of ~43 × 43 μm (x, y) at a 3.5-5 nm per pixel resolution. The imaged volumes were centered to contain the zona pellucida, oocyte surface, and cumulus cell bodies (Figure 1B). We were able to trace every TZP sent by individual cells (Video 1 and Figure 1C), and record their interactions with other TZPs and with the oocyte surface (described below). As can be seen in video 2, TZPs arise from cumulus cells located at varying distances from the oocyte.

We found that the majority of TZPs do not reach the surface of the oocyte (Figure 2A, C, D). On average, each cumulus cell had 31 ± 5 (n = 8) TZPs that do not reach the surface of the oocyte, and 9 ± 2 (n = 8) TZPs that reach the oocyte and make a junction with it (Figure 2B, C, D) (data shown as mean ± standard error of the mean, unless specified otherwise). We will refer to these as free-ended and connected TZPs, respectively.

**Figure 2.**
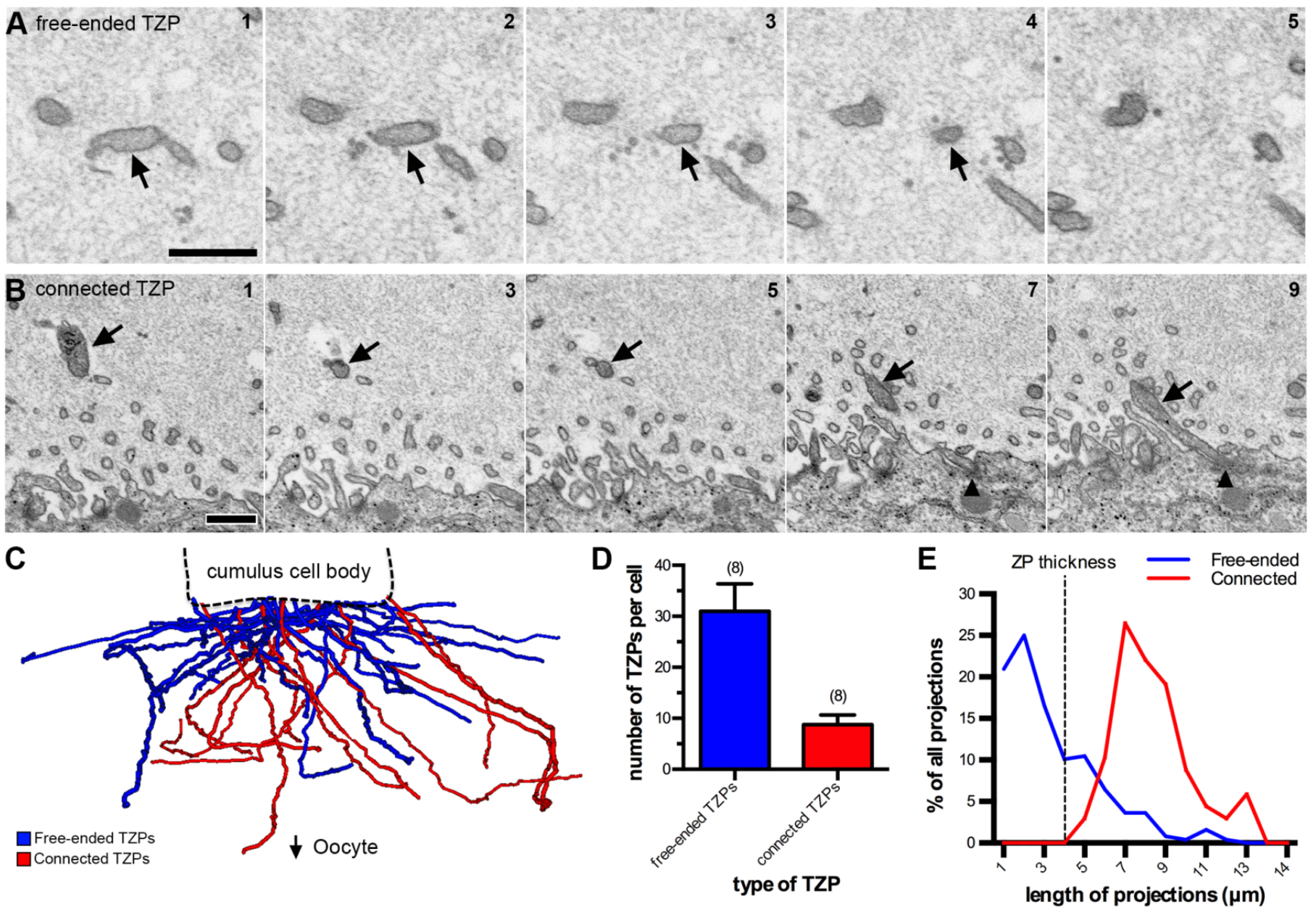
TZPs can be free-ended or connect to the oocyte. **(A)** Serial section SEM images of a free-ended TZP. The TZP (arrow) ends on section 4 without making contact with the oocyte or another TZP. Scale bar, 500 nm. **(B)** A series of SEM images showing every other section of a TZP that connects to the oocyte. Black triangles indicate the TZP-oocyte site of contact. Scale bar, 500 nm. **(C)** Treelines representing every TZP sent by one cumulus cell. Free-ended TZPs are shown in blue, and connected TZPs are shown in red. **(D)** Average number of TZPs per cumulus cell. TZP types were divided into free-ended (blue) and connected (red). 8 cumulus cells from 2 different follicles were analyzed. Total number of projections was 316. Average is shown as mean ± standard error of the mean. **(E)** Histogram of the length of free-ended (blue) and connected (red) TZPs from the data in D. An example of a remarkably long connected TZP can be seen in Video 3.

We measured the lengths of connected and free-ended TZPs in serial electron micrographs (Figure 2E). The lengths of connected TZPs had a bell curve-type distribution with an average length of 7.9 ± 1.9 μm (n = 70) (mean ± standard deviation). Free-ended TZPs had a different kind of distribution, in which the number of projections decreased with length. The first, second, and third quartiles were 1.1, 2.2, and 4.1 μm, respectively. In other words, 25% of the projections were shorter than 1.1 μm, the median (50%) was 2.2 μm, and 25% of the projections were longer than 4.1 μm. The shortest and longest lengths were 0.2 μm and 11.4 μm (see Figure 8C).

Connected TZPs were significantly longer than the average thickness of the zona pellucida, which was 4.1 ± 0.3 μm (n = 3). The thickness of the zona pellucida was measured between the cumulus cell boundary and the oocyte surface at five locations in sections containing the widest diameter of the oocyte. Connected TZPs often extended along the surface of the oocyte making long junctions with it, thereby increasing their average length. Unexpectedly, some connected TZPs looped back to the zona pellucida after making a long junction with the oocyte surface (Video 3). Additionally, many free-ended TZPs were longer than the thickness of the zona pellucida (Figure 2E), but these did not reach the oocyte surface because they traveled obliquely through the zona pellucida (Figure 2C).

Connected TZPs were slightly thicker than free-ended TZPs (75 ± 2 nm (n = 28), and 68 ± 2 nm (n = 30), respectively), and organelles were more often seen in connected TZPs rather than free-ended TZPs. During our analysis, we encountered two cumulus cells in the process of mitosis in two of the follicles analyzed. One cell was in prophase and one in prometaphase. Interestingly, both mitotic cumulus cells had free-ended and connected TZPs (Supplementary Figure 1), suggesting that cumulus cells maintain their connection with the oocyte during mitosis.

### TZPs often contact each other and make gap junctions

Our data revealed novel characteristics of TZPs throughout the zona pellucida. For instance, we found that ~12% of the TZPs analyzed branched into one or more projections (Supplementary Figure 2, and see Figure 3B). Our data also revealed numerous examples of side-to-side and end-to-end contacts between TZPs (Video 4 and Figure 3A). Video 4 shows three examples of TZP contact sites, all found within just 21 sections (a thickness of ~1 μm), providing an example of how common these interactions are within the zona pellucida. Individual contacts like these would be difficult to resolve by light microscopy. TZP-TZP contacts were small, spanning through 1-4 40 nm sections. Most occurred in the upper half of the zona pellucida, closer to the cumulus cells than to the oocyte. We traced interacting TZPs back to their cell of origin to determine if they were derived from different cells (Figure 3B). From 82 TZP-TZP contacts analyzed, 78% were between TZPs from different cells and 22% were between TZPs from the same cell. The presence of same cell TZP-TZP contacts suggests that TZPs form contacts indiscriminately, in other words, two TZPs about to make a contact are not restricted whether they are derived from the same cell or from two different cells.

**Figure 3.**
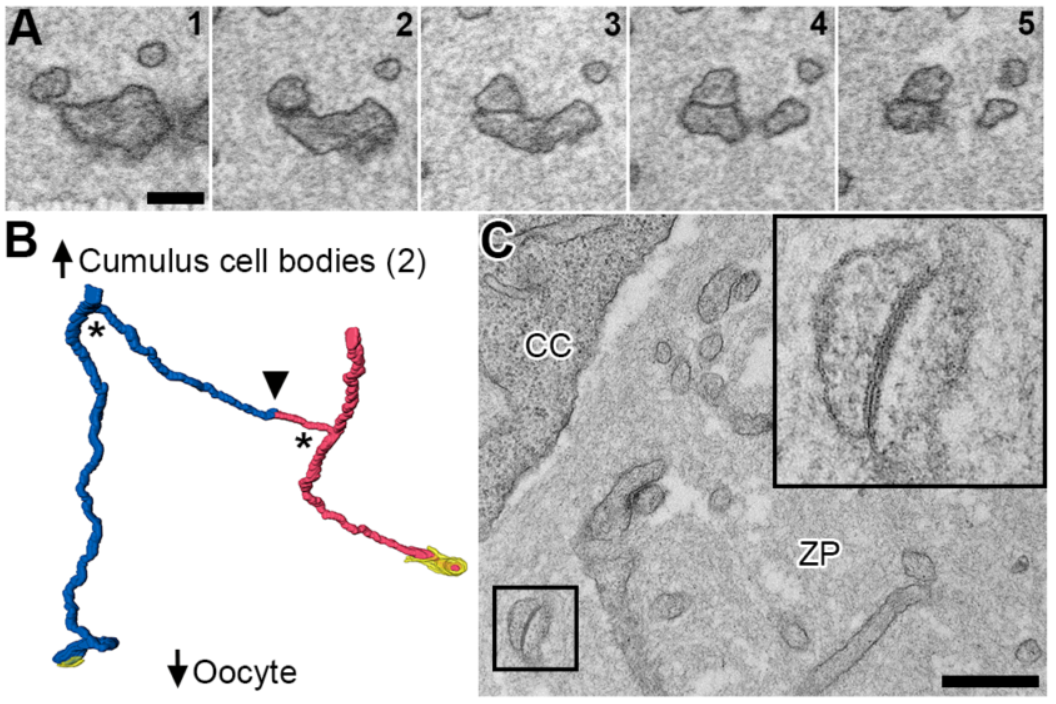
TZPs often contact and make gap junctions with each other. **(A)** Serial section SEM images of two TZPs that contact each other. Scale bar, 250 nm. Video 4 shows three additional examples of contact sites between TZPs. **(B)** Reconstruction of two TZPs derived from different cells that contact each other (black triangle). Asterisks represent branching points on the TZPs (see Supplementary Figure 2). Reconstruction is 4.2 × 4.2 × 5.7 μm (x, y, z), spanning through 126 serial sections (each, 45 nm-thick). Yellow represents the oocyte membrane at the TZP-oocyte junction. **(C)** TEM image of a contact site in the zona pellucida showing a gap junction (high-magnification insert). CC, cumulus cell. ZP, zona pellucida. Scale bar, 500 nm.

Gap junctions within the zona pellucida have previously been observed by immunofluorescence (Simon et al., 2006). However, due to the limited resolution of SEM (~3 nm for these studies), it was not possible to identify small gap junctions in our data. To investigate if the TZP-TZP contacts consisted of gap junctions, we collected sections from the same follicles that had been previously analyzed, and used transmission electron microscopy (TEM) to image the zona pellucidae. We found that most, but not all, contacts in the upper half of the zona pellucida were gap junctions (Figure 3C). Supporting the idea that TZPs can form gap junctions at their endings, we found that some TZPs (18 of 325) ended in an invaginated annular junction within a cumulus cell body (Supplementary Figure 3). Based on our previous immunogold studies (Norris et al., 2017), these are likely to be invaginated gap junctions.

### Connected TZPs and oocyte microvilli make contacts with each other

As described previously (Li and Albertini, 2013; Motta et al., 1994), TZPs end in junctions at the oocyte surface that contain adherens junctions, as identified by an electron-dense deposit (Figure 4A, B) (Niessen and Gottardi, 2008). We reconstructed 12 of these junctions and found that they range from 0.39 μm to 3.59 μm in length, and that most lie along the oocyte surface (Figure 4C), while a few form invaginations into the oocyte cytoplasm.

Mammalian oocytes have a dense network of microvilli on their surface (Runge et al., 2007). In our analysis, oocyte microvilli were generally uniformly distributed along the oocyte surface, and were 1.06 ± 0.09 μm (n = 45) long. Interestingly, we often noticed areas in the zona pellucida where microvilli appeared “clumped” (Figure 4D). Serial section analysis revealed that these clumped areas consisted of one or two TZPs connected to the oocyte, which were closely associated with 3-6 oocyte microvilli (Figure 4E and Video 5). Video 5 shows an example of a connected TZP that is almost continuously coupled with a long microvillus and then becomes surrounded by 5-6 short microvilli as it gets close to the oocyte surface. In some cases, microvilli were seen alongside the TZP for a distance of up to 6 μm (Figure 4F). Although most TZPs that reached the oocyte surface were contacted by microvilli in this manner, some were not. Only one example of a microvillus contacting a free-ended TZP was seen.

**Fig. 4.**
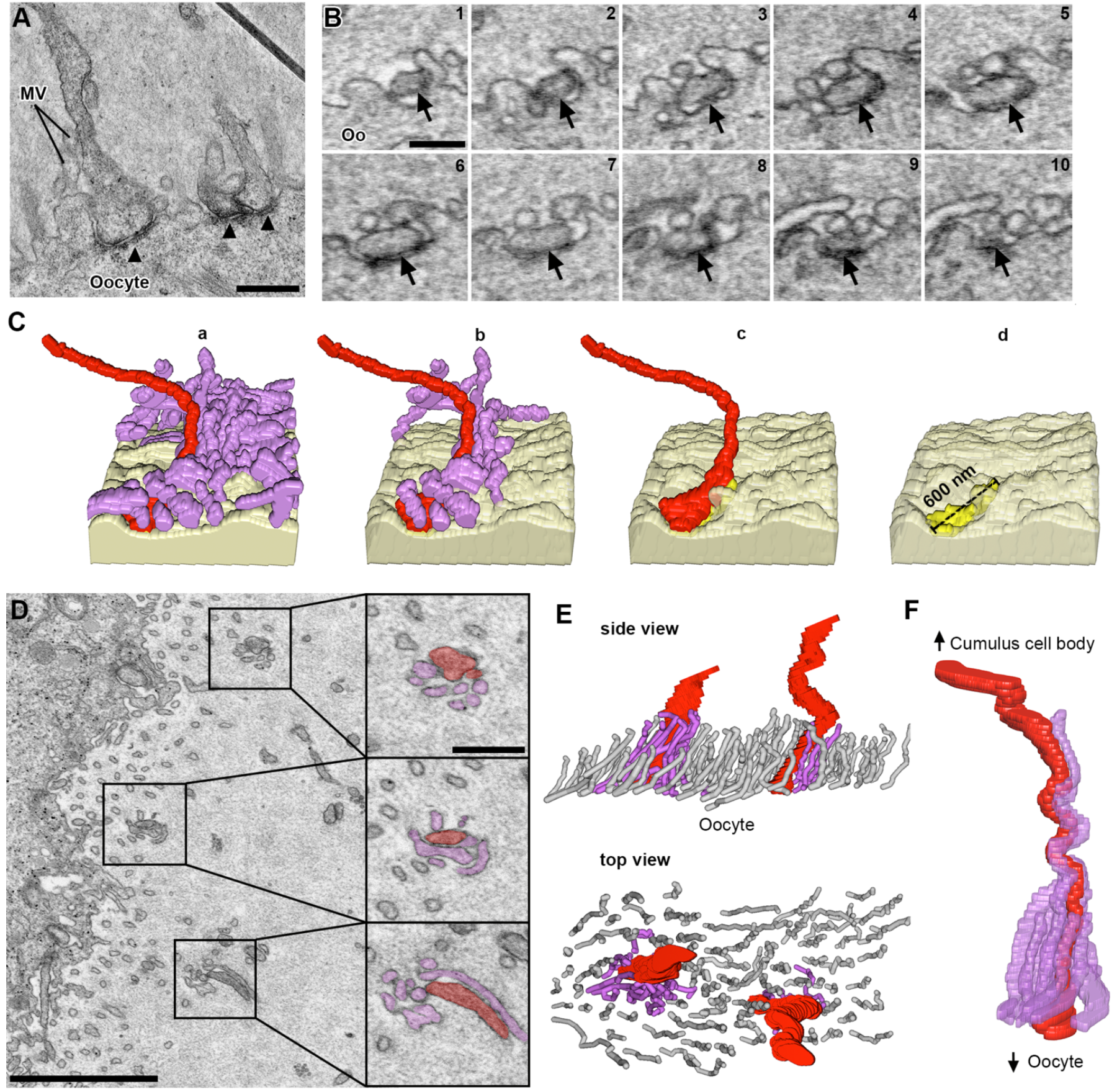
Contacts between TZPs and oocyte components. **(A)** TEM image of TZPs that make adherens junctions with the oocyte surface. Adherens junctions are identified by an electron-dense region at the site of TZP-oocyte contact (black triangles). Mv, oocyte microvilli. Scale bar 500 nm. **(B)** Serial section SEM images of a TZP (black arrow) that makes an adherens junction with the oocyte surface. Oo, oocyte. Scale bar, 300 nm. **(C)** Reconstruction of an adherens junction made by the TZP shown in B. Reconstruction is 2.0 × 1.1 × 1.7 μm (x, y, z), spanning through 38 serial sections (each, 45 nm-thick). Light yellow: oocyte surface. Purple: oocyte microvilli. Red: TZP. Bright yellow: adherens junction. a) TZP and all oocyte microvilli in the volume. b) Unattached microvilli have been removed from the reconstruction to show only those that make a contact with the TZP. c) All microvilli have been removed from the reconstruction. d) TZP has been removed from the reconstruction to show the adherens junction on the oocyte surface. **(D)** SEM image of an area of the zona pellucida in which TZPs and microvilli appear clumped (squares). High-magnification subpanels show the TZP in red and oocyte microvilli in purple (confirmed by serial sections). Scale bars, 2 μm on low magnification, and 500 nm on high magnification subpanels. Video 5 shows this in serial sections (reconstructed in Figure 4F). **(E)** Side and top views of a reconstruction of an area in the zona pellucida that is 3.4 × 1.8 × 2.8 μm (x, y, z), spanning through 63 serial sections (each, 45 nm-thick), showing two areas where TZPs (red) are seen clumped with microvilli from the oocyte (purple). Non-interacting microvilli are shown in gray. **(F)** Reconstruction of a single TZP (red), and oocyte microvilli that tightly contact it (purple). The reconstruction is 2.9 × 1.2 × 2.7 μm (x, y, z), spanning through 61 serial sections (each, 45 nm-thick). This reconstruction was segmented from the data seen in Video 5.

To test whether TZP-microvilli contacts were gap junctions, we inspected thin sections by TEM as described in the previous section. The oocyte cytoplasm and microvilli can usually be distinguished from TZPs by a difference in electron density (see example in Video 3) or by the presence of precipitate that often forms in the oocyte cytoplasm and microvilli, but not on TZPs (see example in Video 5). This allowed us to identify possible TZP-microvilli contacts in the TEM images. In contrast to contacts seen in the upper half of zona pellucida, these did not appear to be gap junctions (data not shown).

### Cytoplasmic projections are also found outside of the zona pellucida (non-TZPs)

During our analysis of TZPs, we found that many cumulus cells had some projections directed away from the oocyte toward other cells, and that mural granulosa cells had similar projections. The number and directionality of the cytoplasmic projections had a striking dependence on cell location within the follicle. Projections not found in the zona pellucida will be referred to as non-TZP cytoplasmic projections.

Cumulus cells were identified as any cell that was connected to the oocyte by means of TZPs. As described before, these cells differed based on whether the cell body was located adjacent to the zona pellucida or displaced away from it (Figure 5A, B). Cumulus cells that were directly adjacent to the zona pellucida extended most projections toward the oocyte as TZPs, and only a few away from it (Figure 5C). These cells had an average of 51 ± 4 TZPs and 10 ± 3 non-TZP cytoplasmic projections (n = 5) (connected and free-ended TZPs were pooled together for these studies). Displaced cumulus cells, which were located 1-2 cell diameters away from the oocyte, had a decreased number of TZPs, and an increased number of non-TZP cytoplasmic projections. These were generally oriented toward other cells and sometimes invaginated into neighboring cell bodies (Figure 5D and Video 1). Video 1 highlights one of these cells and every cytoplasmic projection derived from it, including TZPs. These cumulus cells had 22 ± 9 TZPs and 27 ± 3 non-TZP cytoplasmic projections (n = 3) (see Figure 8A). In summary, cumulus cells that are further displaced from the oocyte have fewer TZPs and more non-TZP cytoplasmic projections compared to cumulus cells adjacent to the oocyte (see Video 2).

**Figure 5.**
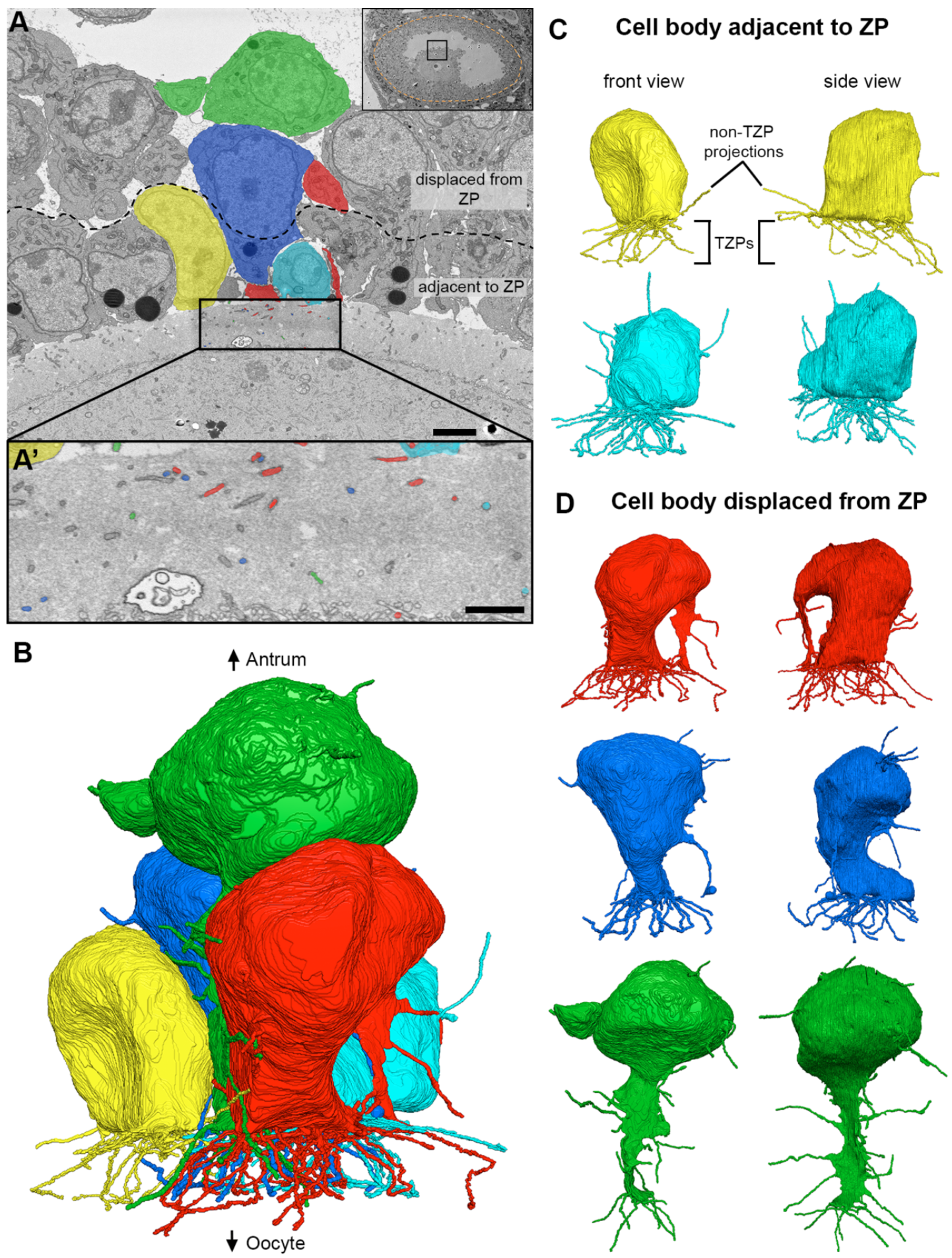
Directionality of cumulus cell projections. **(A)** SEM image showing multiple cumulus cell bodies, zona pellucida, and part of the oocyte in cross-section. Five cells (colored) were chosen for reconstruction. Scale bar, 5 μm. Video 1 shows 202 serial sections of this area and highlights the green cell and all of its projections. **(A’)** High-magnification SEM image of the zona pellucida showing TZPs in cross-section. The colored TZPs originated from the cumulus cells chosen for reconstruction in A. Scale bar, 2 μm. **(B)** Reconstruction of 5 cumulus cell bodies from (A) and every cytoplasmic projection derived from them. Reconstruction is 28.4 × 24.6 × 18.2 μm (x, y, z), encompassing 405 serial sections (each, 45 nm-thick). A rotating view of this reconstruction can be seen in Video 2. **(C-D)** Front and side views of reconstructed cumulus cells found directly adjacent to the zona pellucida (C), or displaced by 1-2 cell diameters (D).

To analyze the projections of mural granulosa cells, we imaged volumes of ~71 × 71 × 27 μm (x, y, z) at a resolution of 5-6 nm per pixel, centered on mural granulosa cells of the follicles that were previously used to study TZPs. Cells chosen for analysis were selected if they were centrally located within the field-of-view and the cell body was completely within the volume.

Inner mural granulosa cells, which are not connected to the oocyte or to the basal lamina (Figure 6A), had an average of 23 ± 2 projections per cell (n = 14). These projections were oriented in many directions with no consistent bias (Figure 6B, C and Videos 6-7). Some of these projections were remarkably long (up to 13.7 μm) and frequently invaginated into neighboring cells. Video 6 shows several examples of such invaginations; in particular, one cell (colored in blue) showed 7 projections, all of which invaginated into its neighboring cell (located to its upper right). We have not detected fused membranes between the invaginated projections and the cells into which they invaginate (See video 10 for an example of an invaginated projection).

**Figure 6.**
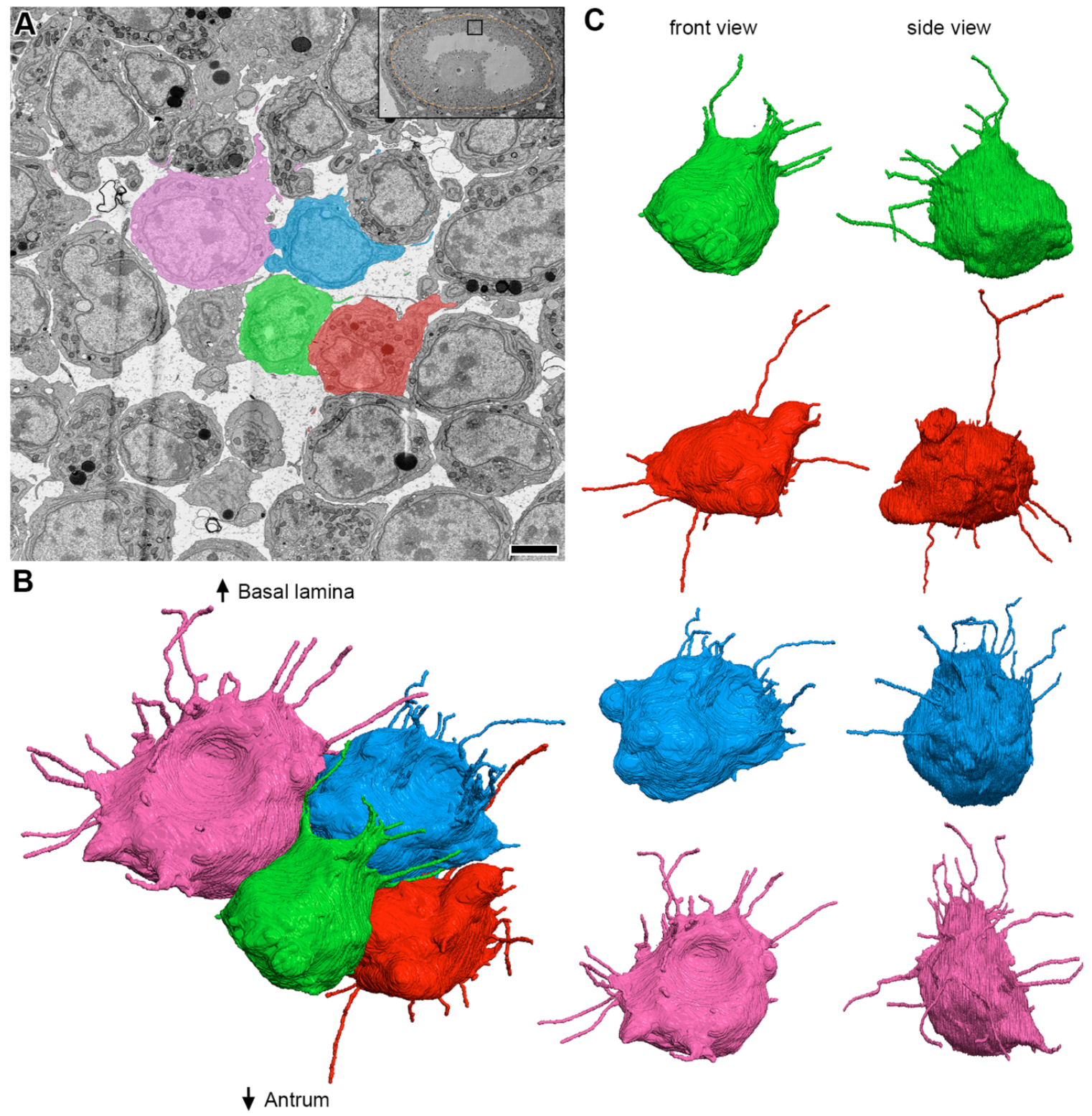
Inner mural granulosa cells send projections in many directions. **(A)** SEM image showing multiple inner mural granulosa cell bodies in cross-section. Four cells (colored) were chosen for reconstruction. Scale bar, 5 μm. Video 6 shows 240 serial sections of these cells and their projections. **(B)** Reconstruction of 4 inner mural granulosa cell bodies from (A) and every cytoplasmic projection derived from them. Reconstruction is 21.2 × 16.4 × 20.8 μm (x, y, z), encompassing 462 serial sections (each, 45 nm-thick). A rotating view of this reconstruction can be seen in video 7. **(C)** Front and side views of individual inner mural granulosa cell reconstructions from (B).

Outer mural granulosa cells were identified as those that had a visible connection with the basal lamina (Figure 7A). Many outer mural granulosa cells had their cell body located 2-3 cell layers away from the basal lamina, but connected to it through a thick long cytoplasmic process (Figure 7B, D) (Lipner and Cross, 1968). Numerous thin cytoplasmic projections, similar to those found in the other cell groups, were seen originating from these elongated cells (Figure 7D and Video 8), with an average of 28 ± 5 projections per cell (n = 4). Strikingly, outer mural granulosa cells that were located directly adjacent to the basal lamina (Figure 7A, C and Video 8) had a significant reduction in the number of projections, having 7 ± 1 (n = 3) per cell. In summary, mural granulosa cells that are not connected to the oocyte or to the basal lamina have numerous projections that are oriented randomly, while cells that are connected to the basal lamina differ based on the location of their cell body: cells that are further displaced from the basal lamina have a larger number of projections compared to cells directly adjacent to the basal lamina (Video 9 and Figure 8A).

**Figure 7.**
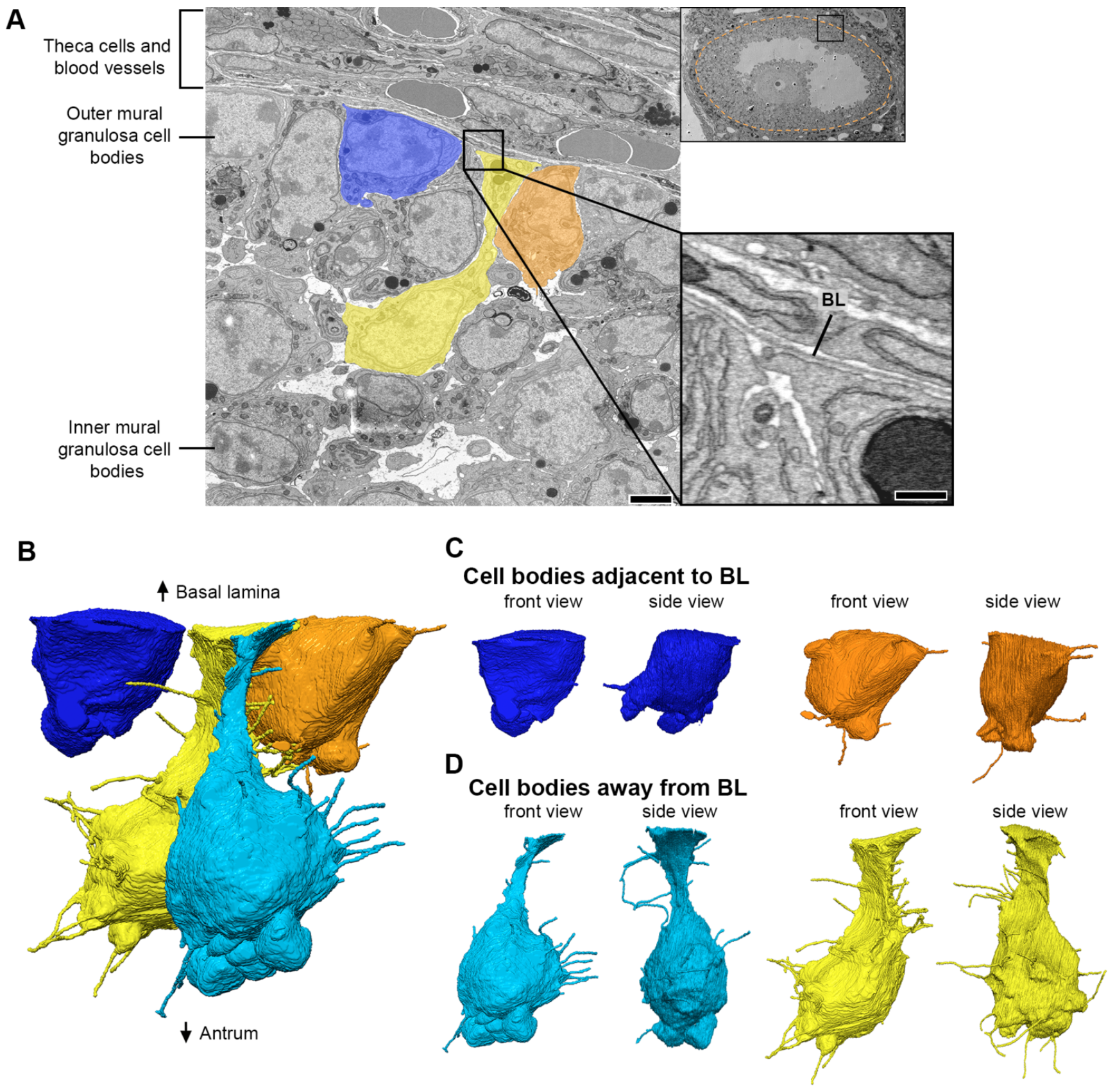
Outer mural granulosa cells send projections in many directions. **(A)** SEM image showing a region of inner and outer mural granulosa cells, the basal lamina, and theca cells and blood vessels found outside of the follicle. Four outer mural granulosa cells were chosen for reconstruction (three colored; one cell was not in the plane of the section). Scale bar, 5 μm. Video 8 shows 267 serial sections of this area. High-magnification insert shows parts of the cell bodies from outer mural granulosa cells on the bottom half and cell processes from theca or endothelial cells on the upper half. The basal lamina (BL) separates these cell types. Scale bar, 500 nm. **(B)** Reconstruction of 4 outer mural granulosa cell bodies and every cytoplasmic projection derived from them. Reconstruction is 16.4 × 14.7 × 22.4 μm (x, y, z), encompassing 497 serial sections (each, 45 nm-thick). A rotating view of this reconstruction can be seen in video 9. **(C-D)** Front and side views of reconstructed outer mural cells from (B), which were located directly adjacent to the basal lamina (C) or 1-2 cell diameters away from it (D).

We then characterized the endings of non-TZP cytoplasmic projections from cumulus, inner mural, and outer mural granulosa cells (analysis of 427 projections from 20 different cells). Most projections (38%) ended on the surface of a neighboring cell. Other common ending types were inside an invagination in a neighboring cell (24%) (Video 10), or as a free end in the extracellular space (26%). Less common endings seen were as an end-to-end contact with a projection from a different cell (7%), as an invaginated annular junction (5%), or as a small linear gap junction (<1%). Examples of all these endings can be seen in videos 1, 6, and 8.

### Common features of projections throughout the follicle

Free-ended TZPs and the cytoplasmic projections of granulosa cells showed a similar overall appearance; both were usually devoid of organelles, and their lengths were strikingly similar. As with free-ended TZPs, we found that the number of non-TZP cytoplasmic projections from the cumulus, inner mural, and outer mural cells decreased with length (Figure 8B). The first, second, and third quartiles were 1.1, 2.5, and 4.4 μm, respectively (described above). The shortest and longest lengths were 0.2 Mm and 13.7 Mm. Figure 8C shows a detailed summary of these findings. The length distribution and thickness of these projections, which was 76 ± 2 nm (n = 67), did not change based on their location within the follicle.

**Figure 8.**
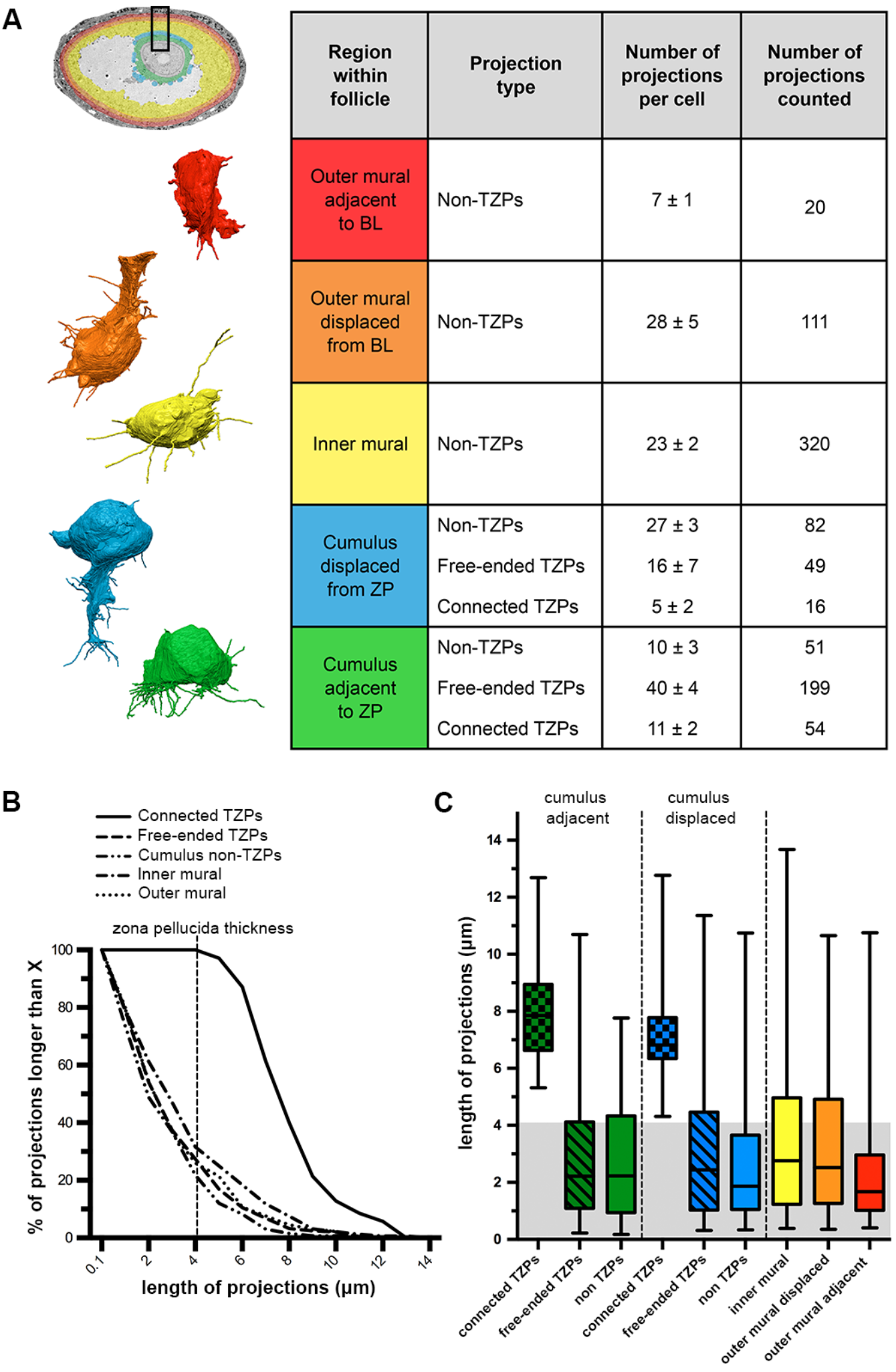
Summary of cytoplasmic projections in somatic cells of antral ovarian follicles. **(A)** Summary table describing the number of projections per cell in each region of the ovarian follicle. Outer mural adjacent to BL refers to cells found directly adjacent to the basal lamina (n = 3). Outer mural displaced from BL refers to cells found two-to-three cell diameters away from the basal lamina but that still connect to it through a thick cytoplasmic process (n = 4). Inner mural refers to cells not connected to the oocyte or to the basal lamina (n = 14). Cumulus displaced from ZP refers to cells found two-to-three cell diameters away from the zona pellucida but that still connect to the oocyte through TZPs (n = 3). Cumulus adjacent to ZP refers to cells found directly adjacent to the zona pellucida (n = 5). Numbers are shown as mean ± standard error of the mean. To the left of the table are representative reconstructions of one cell from each of the cell groups in the table. Follicle insert is colored to represent the different cell groups. **(B)** Distribution of the length of projections from each group. All cumulus cells, regardless of the position of their cell body, were pooled together for the groups free-ended TZPs, connected TZPs, and cumulus non-TZPs. Outer mural granulosa cells were also pooled together. **(C)** Length of projections from every cell group represented as quartiles. Lower and upper edges of the box represent the first and third quartiles, respectively (25^th^ and 75^th^ percentiles). The line in the middle of the box represents the median (50^th^ percentile). The lower and upper limits of the “whiskers” represent the minimum and maximum values, respectively. Cell groups are divided as described in (A). Gray-shaded region represents the thickness of the zona pellucida (~4.1 μm). Note that ~25% of all of the projections from each cell type are longer than the width of the zona pellucida (suggesting they could contact the oocyte if the cell body is at an appropriate distance).

The striking similarity between the lengths of free-ended TZPs and non-TZP cytoplasmic projections from every cell group suggested to us that all granulosa cells are programmed to seek out the oocyte by extending cytoplasmic projections. If so, some of these projections should be longer than the thickness of the zona pellucida. Consistent with this idea, we found that at least a quarter of the projections from every somatic cell within the follicle were longer than the thickness of the zona pellucida, which was 4.1 ± 0.3 Mm (n = 3) (as mentioned above) (Figure 8B, C). Our findings suggest that every somatic cell in the follicle is able to probe a distance that is longer than the thickness of the zona pellucida around their cell body by extending cytoplasmic projections. These projections could allow cells to contact the oocyte if the cell body is located at an appropriate distance (Figure 9).

**Figure 9.**
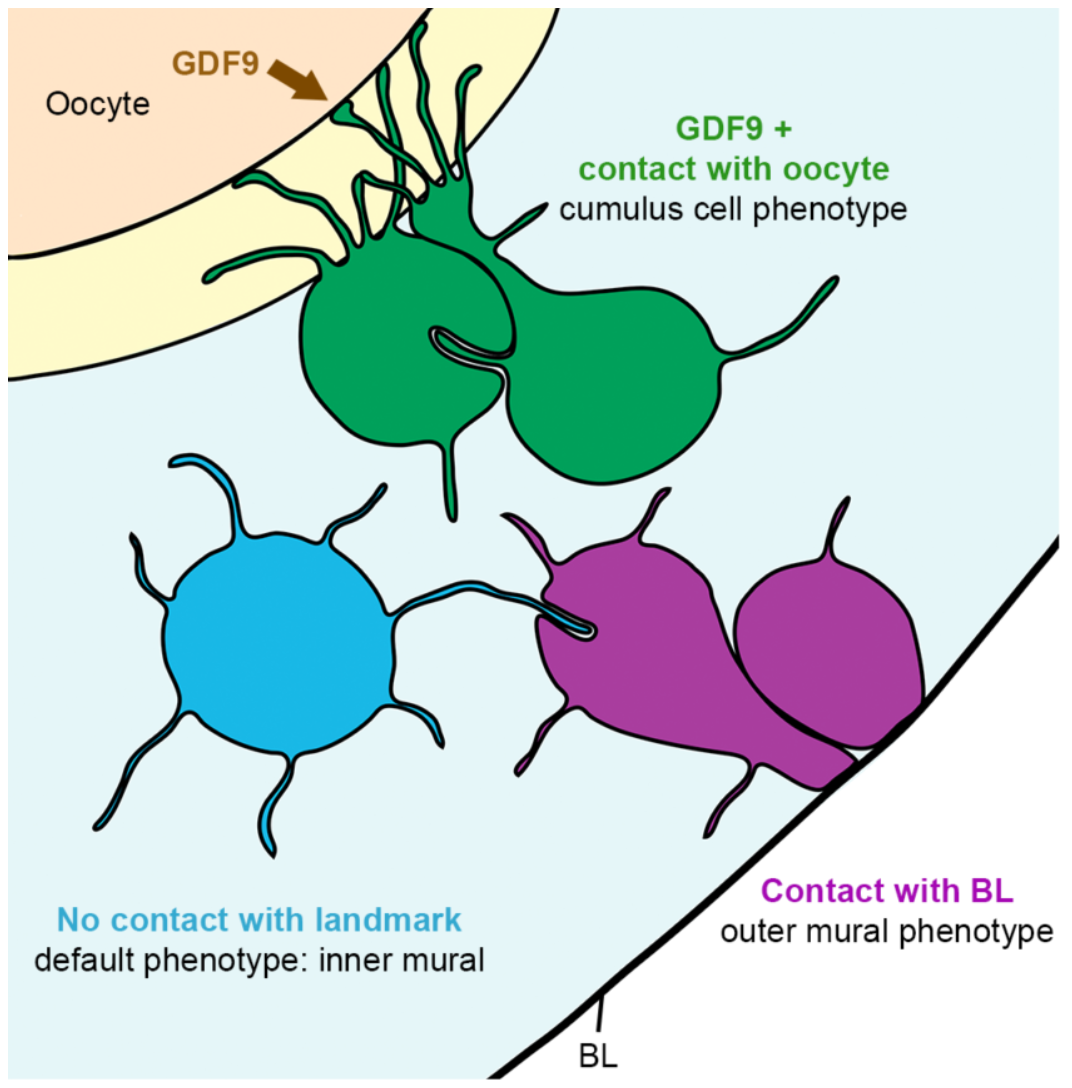
Proposed model. Differentiation of somatic cells in the ovarian follicle is dependent on contact with the oocyte or with the basal lamina. Here, a granulosa cell becomes a cumulus cell (green) if it contacts the oocyte through TZPs and receives the GDF9 signal from the oocyte. If a granulosa cell makes contact with the basal lamina, it becomes an outer mural granulosa cell (purple). Conversely, if a granulosa cell does not make contact with the oocyte or with the basal lamina, it remains as an inner mural granulosa cell (blue), the default phenotype.

## Discussion

In this study, we examined mouse antral ovarian follicles by serial section electron microscopy. Using the ATUM method (Kasthuri et al., 2015), we imaged volumes large enough to examine whole cells at high resolution, along with all of their cytoplasmic projections to determine how they contact other cells in the densely organized regions of the ovarian follicle.

### TZP connections to the oocyte

Previous studies have shown that TZPs contact the oocyte at an adherens junction in a depression in the oocyte surface (Motta et al., 1994). Small gap junctions have also been observed in the vicinity of the oocyte by freeze fracture and thin section TEM (Anderson and Albertini, 1976), and by immunofluorescence (Simon et al., 2006), but these studies did not provide enough three-dimensional information to determine to what degree gap junctions are present at the oocyte surface or on projections from the oocyte.

Our serial section data shows that the TZP lies in a depression in the oocyte surface without much widening (Figure 4C), and can make long linear adherens junctions with the oocyte surface (Video 3).

We found that most TZPs that reach the oocyte surface are contacted by several oocyte microvilli on their shaft (Figure 4D-F and Video 5). This previously undetected association is significant because there are several critical interactions between the oocyte and cumulus cells and one or more of these interactions could occur at these contact sites. We initially considered whether these were the sites of gap junctions between cumulus cells and the oocyte. We tentatively conclude that the microvilli / TZP contact sites are not gap junctions because we did not find them in this region of the zona pellucida by TEM. An interaction that may be occurring at the microvilli / TZP contact sites is discussed below.

### Free-ended TZPs and dynamics

Although TZPs have been known for many years, their dynamics have not yet been characterized. A recent study has focused on this issue using a reconstituted system (El-Hayek et al., 2018). When cumulus cells that have been stripped from their innate oocyte are reaggregated with a donor oocyte, they make new TZPs, which form junctions with the oocyte. This clearly demonstrates that TZPs are dynamic structures.

Our serial section data is consistent with dynamic TZPs in-situ. We show for the first time that most TZPs in the zona pellucida are free-ended (Figure 2). Free-ended TZPs have a wide distribution of lengths and are oriented in many directions. We found that TZPs often make close contacts with each other, some of which were found to be gap junctions by TEM (Figure 3 and Video 4). Our observations are consistent with persistent growth and perhaps retraction of TZPs in intact follicles at a stage when the cumulus cells are already connected to the oocyte by TZPs. All TZPs contain actin but only a small minority contain microtubules (El-Hayek et al., 2018). We suggest that the free-ended TZPs contain only actin and that microtubules grow into them if they connect to the oocyte.

One question is whether the TZP dynamics is induced or constitutive. In their recent study, El-Hayek et al. (2018) present evidence that oocyte-derived factors induce the formation of TZPs. When the oocyte is removed from a granulosa-oocyte complex (GOC), the mRNA levels for general components of filopodia (*Damml*, *Fscnl*, and *Myo10*) are drastically reduced in the remaining granulosa cells. Additionally, when soluble GDF9 (a BMP-like family paracrine factor produced by the oocyte) was added to the GOCs medium, the mRNA levels of filopodia components were restored (there is uncertainty however regarding the physiological dose for GDF9). When GDF9 production was blocked by siRNA injection into oocytes of intact GOCs, filopodia mRNA levels were reduced in granulosa cells and the number of TZPs counted was 30% lower than their controls. The authors conclude that GDF9, possibly with other oocyte-secreted factors, induces cumulus cells to extend filopodia into the zona pellucia towards the oocyte. We present an idea below that the dynamics of TZPs is largely constitutive.

### Cytoplasmic projections / filopodia in the follicle

Our serial section data allowed us for the first time to detect and characterize cytoplasmic projections in parts of the follicle that are densely populated with cells. We found that cumulus cells have some projections directed away from the oocyte (Figure 5 and Video 2), and that mural granulosa cells have numerous projections extending in all directions (Figures 6-7, and Videos 7-9). There is a striking similarity in the length distributions of these projections and the free-ended TZPs (Figure 8). We did not show that these projections contain filopodia markers, but on the basis of their similarity to TZPs, which do have filopodial markers (El-Hayek et al., 2018), it seems likely that these projections are filopodia.

### Possible filopodial functions

Filopodia are frequently seen in cultured cells and are often dynamic when observed by time-lapse imaging. They have also been commonly observed in-situ such as in the sea urchin and Xenopus embryo blastocoel (Miller et al., 1995, Danilchik et al., 2013). There is evidence that filopodia sense environmental cues that guide migration of pathfinding neurons and of vascular endothelial tip cells forming a new capillary (Bentley and Toroian-Raymond, 1986; Fantin et al., 2015). Upon contact with axons, dendritic filopodia were observed to become dendritic spine synapses (Ziv and Smith, 1996).

In the ovarian follicle, filopodia could be involved in transducing paracrine or hormonal signals. These signaling molecules are usually thought to act by diffusion but there is recent evidence that paracrine factors can act by contact with specialized filopodia termed “cytonemes” (Kornberg, 2017). Evidence for signaling by contact has been found in tissues where paracrine signaling is known to occur (Ramirez-Weber and Kornberg, 1999). Receptors and ligands have been localized to filopodia (Sanders et al., 2013; Gonzalez-Mendez et al., 2017), and paracrine signaling is reduced or eliminated by eliminating the filopodia (Roy et al., 2014).

There is evidence for paracrine signaling by contact in the Drosophila male germline stem cell niche. In this system, Decapentaplegic (Dpp), a Drosophila BMP, is secreted by hub cells to maintain adjacent cells as germline stem cells (Kawase et al., 2004). When a germline stem cell divides, the daughter cell, which becomes displaced away from the hub cell (source of Dpp) begins to differentiate. Later work showed that the germline stem cells extend microtubule-based projections (MT-nanotubes), which invaginate into hub cells. Dpp and its receptor Thickveins (Tkv) were localized to these invaginated compartments (Inaba et al., 2015). Thus, the Dpp / Tkv interaction occurs where two cells make a close contact.

We attempt here to provide an explanation for our observations of filopodia in granulosa cells and the TZPs in cumulus cells. In the follicle, it is thought that the inner mural granulosa cell is the default state and that GDF9 induces the cumulus cell-specific phenotype. Cultured granulosa cells from dissociated follicles are induced to synthesize cumulus cell markers by addition of soluble GDF9 (Elvin et al, 1999) and a partial knockdown of GDF9 causes diminished expression of cumulus cell markers (Su et al., 2004).

It is assumed that GDF9 diffuses from the oocyte across the zona pellucida and induces adjacent cells to become cumulus cells. We propose instead that the GDF9 interactions occur when a filopodia contacts the oocyte. In this idea, granulosa cells use filopodia to search whether they are close to the oocyte. If they contact the oocyte, they become a cumulus cell, and the contacting filopodia convert into a connected TZP. Conversely, if a granulosa cell is in contact with the basal lamina, it becomes an outer mural granulosa cell (Figure 9).

Our proposal accounts for the presence of filopodia throughout the follicle as well as their similar length distributions, in which ~25% are long enough to traverse the zona pellucida (Figure 8C). The main prediction is that receptors for GDF9 are present on the filopodia. In particular, these receptors should be present at sites of contact of oocyte and TZPs, for instance, at the microvilli / TZP contact sites. Receptors for GDF9 have been localized but at a resolution too low to address this issue (Sun et al., 2010). New methods for immunolabeling serial sections may help to test this idea (Norris et al., 2017).

In summary, new methods for serial section electron microscopy enabled us to examine whole cells and their relationship to other cells with ultrastructural resolution. Our study shows that this kind of structural information may be useful in understanding the interactions that occur during development and function of a complex tissue structure. Knowledge of other tissue structures is likely to benefit from similar serial section analysis.

## Materials and methods

### Animals

Ovaries were obtained from prepubertal 24-day-old C57BL/6J mice (Jackson Laboratories, Bar Harbor, ME). The mice were not injected with hormones prior to euthanasia. All procedures were approved by the animal care committee at UConn Health.

### Tissue processing for electron microscopy

After dissection from the animal, ovaries were cleaned and split into 3-4 pieces with forceps. Special attention was used to attempt to preserve follicle integrity. The pieces were rinsed in 1× PBS once, and then fixed in a Karnovsky’s fixative (2.5% glutaraldehyde / 2% paraformaldehyde) in 0.1 M cacodylate buffer for 3-4 hours at room temperature. Ovaries were then rinsed several times in 0.1 M cacodylate buffer and stored overnight at 4°C.

Ovaries were post-fixed with 4% OsO_4_ in 3.2% potassium ferricyanide in 0.1 M cacodylate buffer for 1 hour at room temperature. They were then thoroughly rinsed with distilled water, treated with 1% aqueous uranyl acetate overnight at 4°C, and then treated with 0.066% lead aspartate for 30 minutes at 60°C. The samples were then rinsed thoroughly with distilled water, dehydrated in graded ethanol solutions, embedded in epoxy resin (Poly/bed 812 Embedding Media, Polysciences, Warrington, PA), and polymerized in a 60°C oven for 48 hours.

### Sectioning, imaging, and analysis

Serial sections were cut from the polymerized epon blocks with a diamond knife (Ultra 45°, Diatome, Hatfield, PA) at a thickness of 40-45 nm. For each sample, 400-600 serial sections were collected on tape using an automated tape-collecting ultramicrotome (Kasthuri et al., 2015). The tape with sections was laid on silicon wafers (University Wafer, Boston, MA), and then coated with carbon.

The sections were mapped and imaged as described previously (Terasaki et al., 2013) using a field-emission scanning electron microscope (Zeiss Sigma FE-SEM) in backscatter mode (8 keV), at a resolution of 3.5-6 nm/pixel (12,000 × 12,000 pixels), and the Atlas-4 Imaging software (Fibics, Ottawa, Ontario, Canada) in conjunction with custom scripts. Some sections were additionally imaged with a Verios 460L field-emission scanning electron microscope (FEI, Raleigh, NC). These were taken by the backscatter detector (5 keV) using immersion mode, at a resolution of 3-10 nm/pixel.

For TEM imaging, blocks that had already been cut for SEM serial sections were re-trimmed into a ~1 mm × 1 mm face, and 4-6 60 nm serial sections were collected as ribbons on formvar-coated slot grids. Sections were additionally post-stained with 3% aqueous uranyl acetate for 5 minutes, and dried at room temperature. Grids were imaged with a Hitachi H-7650 transmission electron microscope (Hitachi, Tarrytown, NY).

FIJI Image J was used for the image analysis. The alignment was done using the Linear Stack Alignment with SIFT plugin. The segmentations were done in the TrackEM2 module (Cardona et al., 2012) using area lists, and the measurements of cytoplasmic projections were done in the same module using treelines. The reconstructions were rendered in Adobe Photoshop CS6-Extended (Adobe, San Jose, CA).

## Acknowledgements

We thank Arthur Hand and Maya Yankova for providing training and suggestions on processing and sectioning samples for EM, Richard Schalek and Jeff Lichtman for advice and continuous support on serial section EM, Ninna Shuhaibar for helping with segmentations, and Rindy Jaffe, Rachael Norris, and Mayu Inaba for thoughtful discussions and critical review of the manuscript. This work was supported by a grant from the Connecticut Science Fund.

---

**Supplementary Figure 1.**
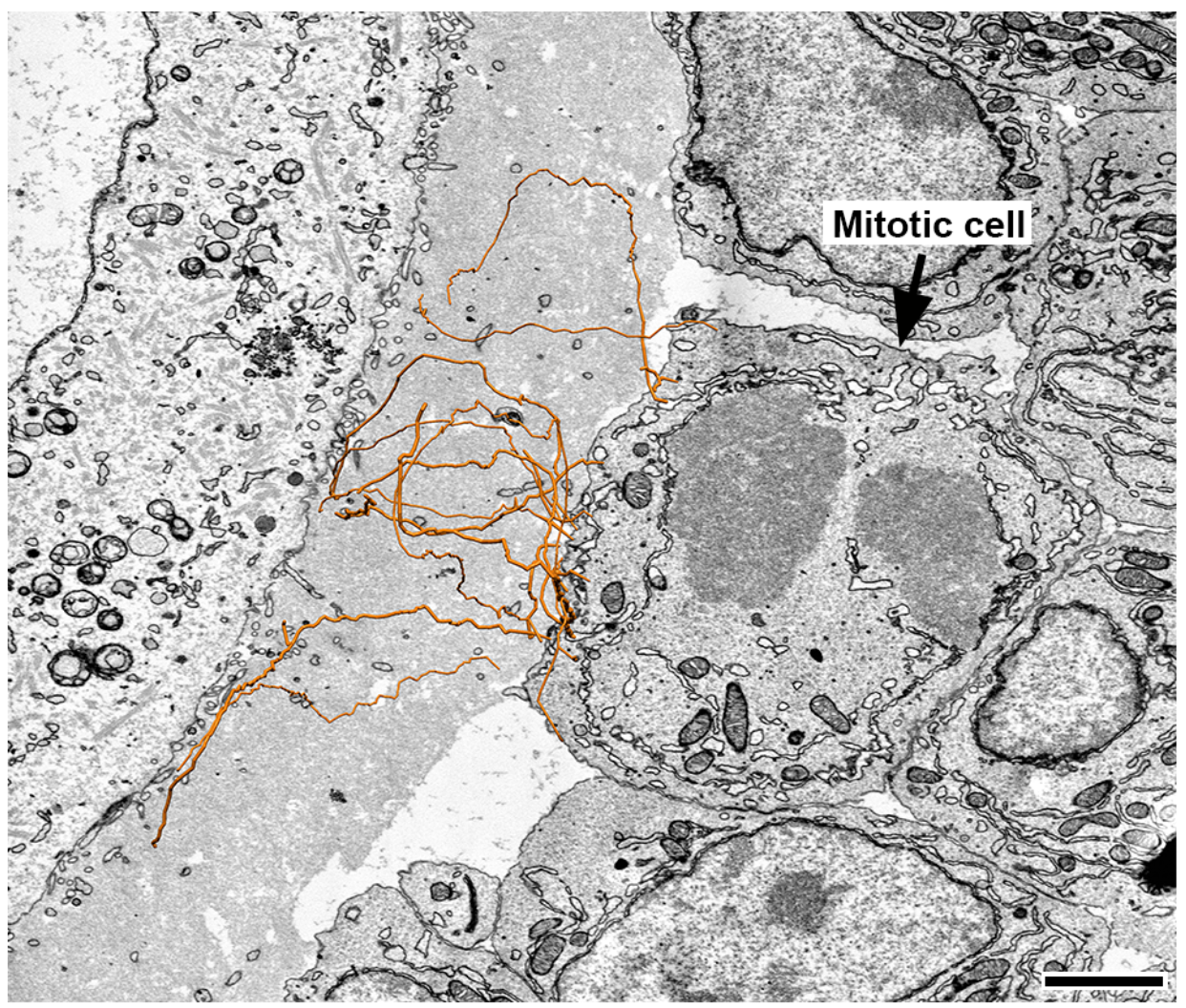
Mitotic cumulus cells have connected and free-ended TZPs. Mitotic cumulus cells were identified by the presence of condensed chromatin and the absence of a nuclear envelope. TZPs derived from this cell were reconstructed from serial sections and then overlaid on the micrograph (orange).

**Supplementary Figure 2.**
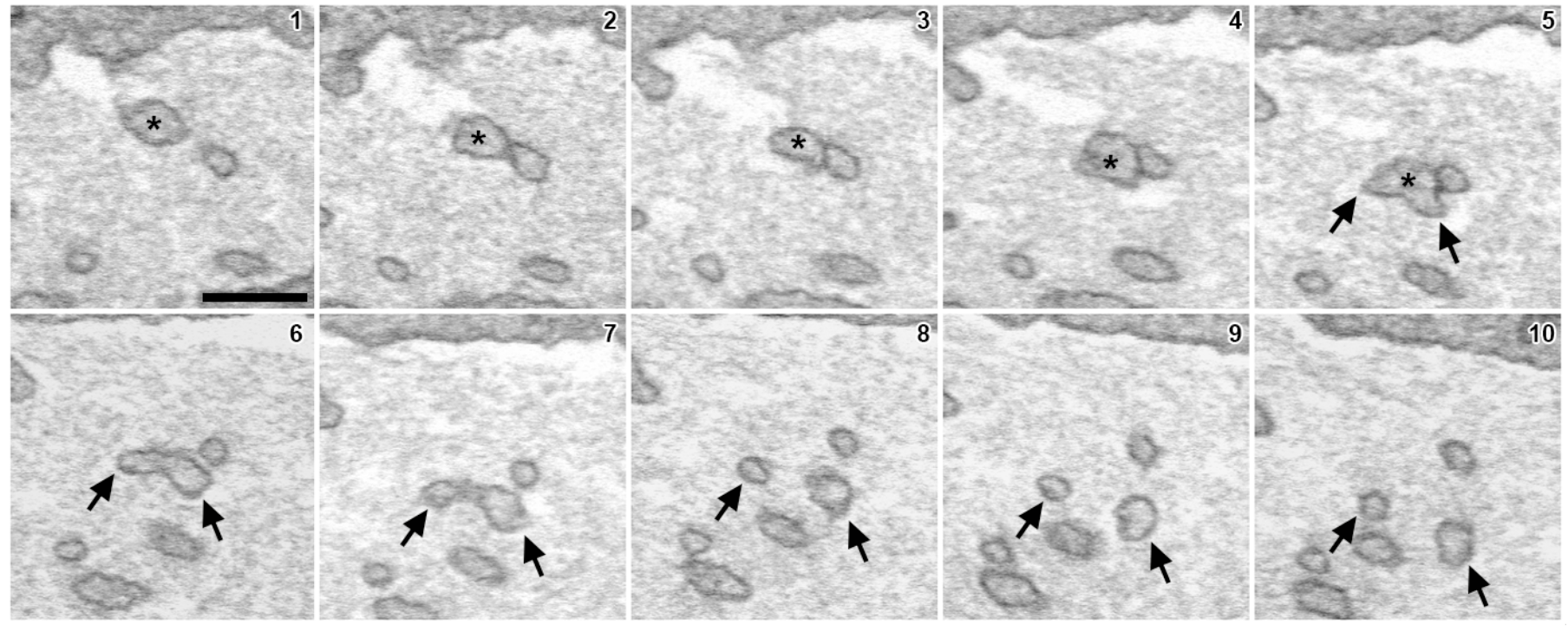
TZP branching. A TZP branches into two projections. An asterisk labels the original TZP. Two black arrows label each of the TZP branches. Figure 3B shows a reconstruction of TZPs in which two branching points can be seen.

**Supplementary Figure 3.**
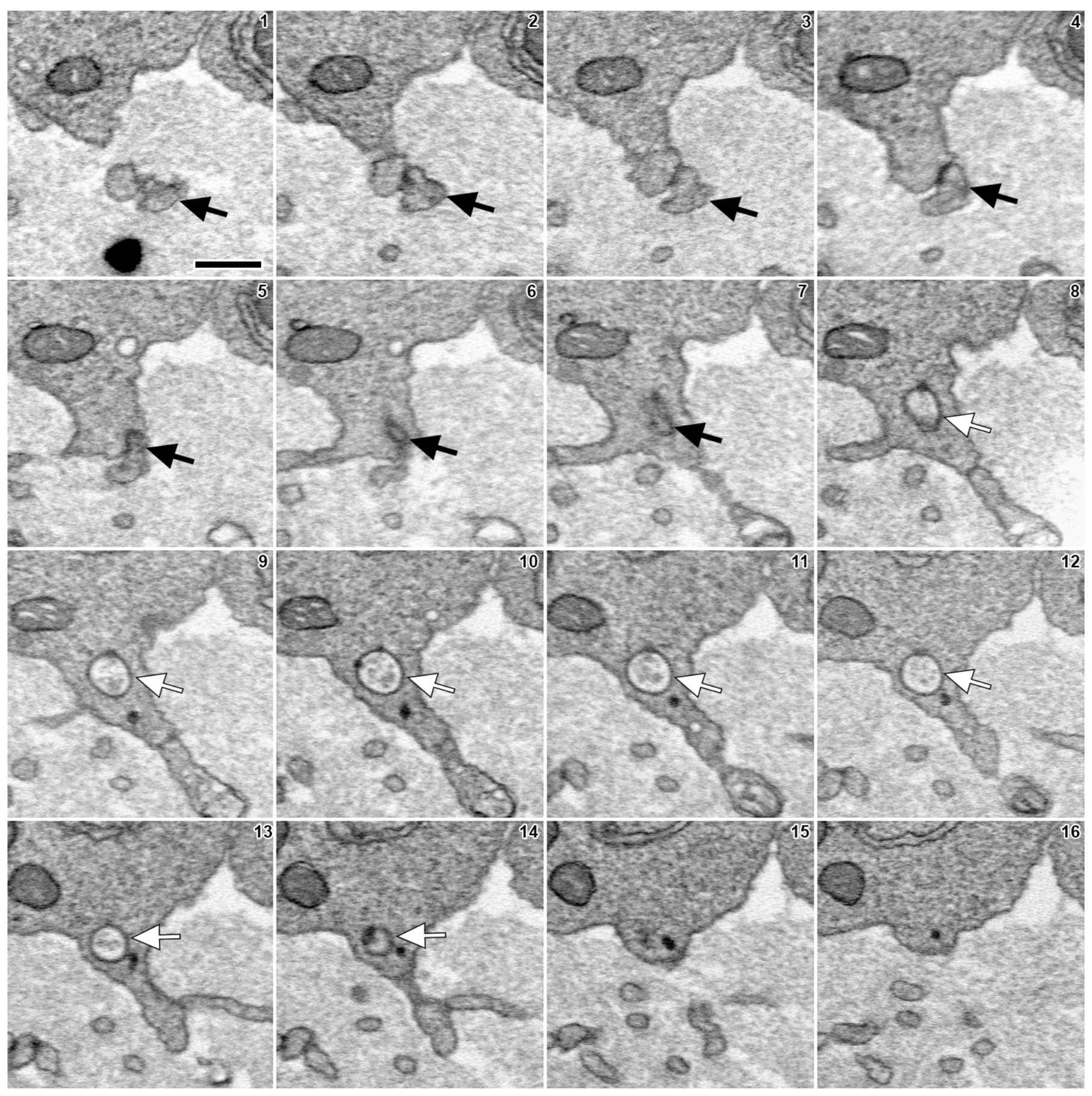
TZPs make invaginated junctions with cumulus cell bodies. Black arrow labels a TZP that invaginates the cell body of a cumulus cell. Based on comparison with our previous study, this is most likely an invaginated gap junction (Norris et al., 2017).

## Figure legends for supplementary videos

**Video 1. Projections can be derived from cumulus cells not directly adjacent to the zona pellucida.** 202 serial section SEM images highlighting a cumulus cell that is displaced from the zona pellucida. Connected and free-ended TZPs originate from the cell process at the edge of the zona pellucida. Additionally, several cytoplasmic projections originate from the elongated shaft of the cell and travel in many directions. All cytoplasmic projections originating from this cell body are labeled in green. Same cell shown in Figure 5D (green).

**Video 2. Reconstruction of 5 cumulus cell bodies and every cytoplasmic projection derived from them.** Reconstruction is 28.4 × 24.6 × 18.2 μm (x, y, z), encompassing 405 serial sections (each, 45 nm-thick). The oocyte and the zona pellucida are located at the bottom. Same reconstruction as Figure 5B.

**Video 3. A connected TZP makes a long looping junction with the oocyte surface.** Serial section SEM images through the zona pellucida of an antral follicle. Cumulus cell bodies are shown on top, and the oocyte surface is shown on bottom. Asterisk labels a connected TZP that makes a long junction with the oocyte surface and then loops back into the zona pellucida.

**Video 4. TZPs make contact sites with each other.** Serial electron micrographs showing three examples of contact sites between TZPs. Each contact site is labeled with a differently colored arrow. Most of these contacts were found to be gap junctions by TEM (see Figure 3C).

**Video 5. Oocyte microvilli closely associate with connected TZPs (serial sections).** Red arrow labels a TZP that eventually connects to the oocyte surface. Yellow arrow labels a long microvillus from the oocyte that associates with the TZP for a long distance. The TZP gets surrounded by more microvilli (intermittent yellow arrows) as it gets closer to the oocyte surface. A reconstruction of these structures can be seen in Figure 4F.

**Video 6. Inner mural granulosa cells send projections in many directions (serial sections).** 240 serial section SEM images of inner mural granulosa cells. Four cells (same as those in figure 6) are labeled with different colors. Cytoplasmic projections derived from each cell are labeled with the same color as their respective cell body. Several long projections that extend for several cell diameters can be seen originating from every cell. Notice multiple projections that originate from the blue cell and invaginate extensively into a neighboring cell located to its upper right. A rotating reconstruction of these cells can be seen in video 7.

**Video 7. Inner mural granulosa cells send projections in many directions (reconstruction).** Reconstruction of 4 inner mural granulosa cell bodies and every cytoplasmic projection derived from them. Reconstruction is 21.2 × 16.4 × 20.8 μm (x, y, z), encompassing 462 serial sections (each, 45 nm-thick). Same cells shown in Figure 6 and Video 6.

**Video 8. Outer mural granulosa cells send projections in many directions (serial sections).** 267 serial section SEM images of outer mural granulosa cells. Four cells (same as those in figure 7) are labeled with different colors. Cytoplasmic projections derived from each cell are labeled with the same color as their respective cell body. The basal lamina is shown at the top and cell processes from theca and endothelial cells are seen on the opposite side of it (see Figure 7A). Notice that the yellow and light blue cells connect to the basal lamina through a long thick cytoplasmic process. Thin cytoplasmic projections can be seen originating from the cell bodies and the thick cytoplasmic process. Numerous invaginating projections originate from the yellow cell towards a neighboring cell to its right. A rotating reconstruction of these cells can be seen in video 9.

**Video 9. Outer mural granulosa cells send projections in many directions (reconstruction).** Reconstruction of 4 outer mural granulosa cell bodies and every cytoplasmic projection derived from them. Basal lamina is located at the top. Notice that the further away the cell body is from the basal lamina, the more projections it possesses. Reconstruction is 16.4 × 14.7 × 22.4 μm (x, y, z), encompassing 497 serial sections (each, 45 nm-thick). Same cells shown in Figure 7 and Video 8.

**Video 10. Non-TZP cytoplasmic projections often invaginate into neighboring cells.** Red arrow labels a cytoplasmic projection derived from a cumulus cell, which invaginates into a neighboring cumulus cell. Notice the space between the invaginated projection and the plasma membrane of the cell into which it invaginates. We have not detected fused membranes at the invaginations. This type of ending is seen in 24% of all non-TZP cytoplasmic projections of every cell type.

